# Random clonal expansion as a limiting factor in transplantable *in vivo* CRISPR/Cas9 screens

**DOI:** 10.1101/2021.11.28.469740

**Authors:** Tet Woo Lee, Francis W. Hunter, William R. Wilson, Stephen M.F. Jamieson

**Affiliations:** Auckland Cancer Society Research Centre, University of Auckland, Auckland, New Zealand; Maurice Wilkins Centre for Molecular Biodiscovery, University of Auckland, Auckland, New Zealand; Discovery Sciences, Janssen Research and Development, Spring House, PA, USA; Department of Pharmacology and Clinical Pharmacology, University of Auckland, Auckland, New Zealand

## Abstract

Transplantable *in vivo* CRISPR/Cas9 knockout screens, in which cells are transduced in vitro and inoculated into mice to form tumours in vivo, offer the opportunity to evaluate gene function in a cancer model that incorporates the multicellular interactions of the tumour microenvironment. In this study, we sought to develop a head and neck squamous cell carcinoma (HNSCC) tumour xenograft model for whole-genome screens that could maintain high gRNA representation during tumour initiation and progression. To achieve this, we sought early-passage HNSCC cell lines with a high frequency of tumour initiation-cells, and identified the pseudodiploid UT-SCC-54C line as a suitable model from 23 HNSCC lines tested based on a low tumourigenic dose for 50% takes (TD_50_) of 1100 cells in NSG mice. On transduction with the GeCKOv2 whole-genome gRNA library (119,461 unique gRNAs), high (80-95%) gRNA representation was maintained in early (up to 14 d) UT-SCC-54C tumours in NSG mice, but not in UT-SCC-74B tumours (TD_50_=9200). However, loss of gRNA representation was observed in UT-SCC-54C tumours following growth for 38-43 days, which correlated with a large increase in bias among gRNA read counts due to stochastic expansion of clones in the tumours. Applying binomial thinning simulations revealed that the UT-SCC-54C model would have 40-90% statistical power to detect drug sensitivity genes with log_2_ fold change effect sizes of 1-2 in early tumours with gRNA libraries of up to 10,000 gRNAs and modest group sizes of 5 tumours. In large tumours, this model would have had 45% power to detect log_2_ fold change effect sizes of 2-3 with libraries of 2,000 gRNAs and 14 tumours per group. Based on our findings, we conclude that gRNA library size, sample size and tumour size are all parameters that can be individually optimised to ensure transplantable *in vivo* CRISPR screens can successfully evaluate gene function.

## Introduction

The molecular profiling of human tumours offers unprecedented opportunities to individualise cancer therapy (Tsimberidou, Eggermont, and Schilsky 2014).

Detection of oncogenic mutations in patient tumours allows treatment with therapies that directly target the mutation, in some instances in a tissue-agnostic approach (Dittrich 2020). However, the role of gene expression in patient response to therapy is less clear. Gene expression profiles can be used to track emergence of resistance (Sun et al. 2014; Murtaza et al. 2013) and to identify predictive biomarkers to optimally match drugs with patients (McKeage et al. 2013), but there is an incomplete understanding of the genes that determine sensitivity to most anticancer agents. One approach to remedy this is through the use of functional genomics screens, which utilise gene knockdown (RNA interference) or gene knockout (CRISPR/Cas9) to select specific phenotypes (e.g. drug sensitivity or resistance) from libraries of mutants and then identify the gene mutation responsible (Hunter et al. 2017).

Whole genome functional genomics screens utilising CRISPR/Cas9 technology were first applied for evaluating genes involved in drug sensitivity or resistance in cancer cell lines in 2014 (T. Wang et al. 2014; Shalem et al. 2014; Zhou et al. 2014). Since that time, a large number of in vitro whole genome screens have been undertaken utilising multiple different guide RNA (gRNA) libraries to identify essential genes (Hart et al. 2015; T. Wang et al. 2015; 2017) and uncover novel genes implicated in tumour growth (Behan et al. 2019; Meyers et al. 2017), drug sensitivity and resistance (Kurata et al. 2016; Bester et al. 2018; MacLeod et al. 2019; Wei et al. 2019; Pettitt et al. 2018), synthetic lethal interactions (T. Wang et al. 2017; C. Wang et al. 2018; Dhoonmoon et al. 2020; Steinhart et al. 2017) and immunotherapy response (Patel et al. 2017; Burr et al. 2017). However, while CRISPR/Cas9 screens provide a powerful approach for gene discovery in vitro, the complex multicellular interactions within the tumour microenvironment (Junttila and De Sauvage 2013; Singleton, Macann, and Wilson 2021) means that the genes controlling cell growth and drug sensitivity are not necessarily the same in cell culture and in tumours. To address this, tumour models are required that are suitable for *in vivo* functional genomics screens.

Tumour models for *in vivo* CRISPR screens can be generated through direct mutagenesis *in vivo* or through *in vivo* transplantation of cells mutagenised in culture. The direct approach allows spontaneous autochthonous tumour formation with preservation of the tumour microenvironment in immunocompetent mice, but is associated with numerous technical challenges, restricting its use to small gRNA libraries (Chow et al. 2017; G. Wang et al. 2018) . Transplantable models, conversely, are more amenable to larger gRNA libraries. Cancer cell lines transduced at scale with whole-genome gRNA libraries can be injected into mice at large cell numbers to ensure high representation of each gRNA at inoculation (S. Chen et al. 2015; Song et al. 2017; Yau et al. 2017; Rushworth et al. 2020; Kodama et al. 2017; Dai et al. 2021). However, the majority of whole-genome transplantable *in vivo* CRISPR screens report a dramatic loss of clonal diversity during tumour growth, due to the selection and bottleneck of cellular evolution imposed by the transition from in vitro to *in vivo* tumour growth (S. Chen et al. 2015; Song et al. 2017; Rushworth et al. 2020; Kodama et al. 2017). This loss of clonal diversity and gRNA representation consequently limits the use of whole genome *in vivo* CRISPR screens for the discovery of therapeutic genes.

Intratumour heterogeneity drives tumour evolution as clonal subpopulations adapt differently to selective pressures through Darwinian selection (Gerlinger et al. 2012; McGranahan and Swanton 2017; Nik-Zainal et al. 2012). In xenografts, a recent single-cell lineage tracing study revealed metastasis is driven by clones with high expression of metastasis-associated genes prior to implantation in mice (Quinn et al. 2021), suggesting Darwinian selection also applies during tumour initiation in xenografts. Yet, little is known regarding the genetic bottleneck when large numbers of cells are transplanted into mice. In this study, we evaluate the tumour initiating ability of early-passage head and neck squamous cell carcinoma cell lines and develop a tumour xenograft model for *in vivo* CRISPR/Cas9 cell transplantation screens from a head and neck squamous cell carcinoma cancer cell line with a high proportion of tumour-initiating cells. We use this tumour model for whole-genome screens using a large gRNA library to evaluate if the genetic bottleneck in tumour initiation and/or progression reflects stochastic effects due to a small number of tumour-initiating cells acting as founders. We also perform simulations using binomial thinning to determine statistical power to detect drug sensitivity genes in small and large tumours with differing gRNA library sizes to provide a demonstration of the key parameters that must be optimised to ensure successful *in vivo* CRISPR screens.

## Methods

### Lentiviral packaging

lentiCas9-Blast (AddGene 52962) was packaged in HEK293T cells using the pMD2.G (Addgene 12259) and psPAX2 (Addgene 12260) packaging plasmids transfected with Lipofectamine 2,000 (Thermo Fisher Scientific). The Human GeCKOv2 A and B libraries in lentiGuide-Puro were amplified as described (Joung et al. 2017), pooled at equimolar concentrations and packaged in the same way. Filtered unconcentrated lentiCas9-Blast lentivirus was used for transductions, while GeCKO library lentivirus was concentrated by centrifugation at 10,000 × g for 5 h with a 20% sucrose cushion.

### Transductions

To produce Cas9-expressing UT-SCC-54C and UT-SCC-74B cells, 5 × 10^4^ cells in normal culture medium with 8 µg/ml polybrene were transduced with lentiCas9-Blast lentivirus at a multiplicity of infection (MOI) of 0.2-0.4. Blasticidin (10 µg/ml) selection was applied 24 h after transduction to select for stable transductants. Cas9-expressing cells were expanded and transduced at an MOI of 0.2-0.3 to produce UT-SCC-54C and 74B GeCKO cell populations (UT-SCC-54C/GeCKO and UT-SCC-74B/GeCKO). Puromycin selection (1-1.5 µg/ml) was applied 2 days after transduction and the proportion of resistant cells was determined after 2 additional days. The MOI was determined using the proportion of resistant cells and Poisson statistics. A total of 6.4 × 10⁷ cells UT-SCC-54C were transduced with a MOI of 0.22 to obtain 1.25 × 10⁷ individual transductants (estimated 89% receiving a single gRNA), and for UT-SCC-74B, 6 × 10⁷ cells were transduced with a MOI of 0.18 to obtain 9.8 × 10⁶ individual transductants (estimated 91% receiving a single gRNA). Cell populations were expanded and either cryopreserved at scale (20-30M cells/tube) or used immediately for inoculations. An early timepoint sample of transduced cells (50M) was obtained at the time zero (T0) sample.

### Animal experiments

Six to eight week old female NOD scid gamma (NSG; NOD.Cg-Prkdc^scid^ Il2rγ^tm1Wjl^/SzJ; Jackson Laboratory) or NIH-III mice were inoculated with UT-SCC cells subcutaneously into bilateral flanks. All animals had ad libitum access to food and water in microisolator cages and were maintained on a 12 h light/dark cycle. Animal health and welfare was monitored regularly with animals culled if their condition deteriorated or if they lost in excess of 20% of their pre-manipulation bodyweight. Tumour volume was recorded by electronic calipers using the formula π/6 x width x length^2^. A tumour was considered to have formed if it reached 250 mm³ in volume within 100 days of inoculation. For TD_50_ determinations, we initially screened all cell lines for growth as tumour xenograft models in NIH-III mice by inoculating 5 × 10⁶ cells per flank into 3-5 mice, followed by 10⁶ and 10^5^ cells for those with a high take rate (>75%). For tumour models that grew as xenografts at 10^5^ or 10⁶ cells, we proceeded to test in NSG mice, which were expected to be more conducive to tumour growth, at 10⁵, 10⁴ and 10³ cells to determine the TD_50_ for each cell line. TD_50_ values were determined using logistic regression fitted to the proportion of tumours that developed (Meyer, Hopwood, and Gillette 1978). For GeCKO cell populations, 10⁷ cells were inoculated. All animal experiments followed procedures approved by the University of Auckland Animal Ethics Committee (approvals #001190 and #001781).

### Tumour collection and drug treatment

In initial studies with UT-SCC-54C/GeCKO and UT-SCC-74B/GeCKO cells, tumours were harvested once the smallest tumour in each group was ∼250 mm³ or greater. For follow-up experiments with UT-SCC-54C/GeCKO, 10⁷ cells were inoculated bilaterally and grown until the median tumour size reached ∼250 mm³, and then mice were randomly divided into four groups of 4-5 mice per group: For group A (n=4), mice were culled and tumours were collected, group B (n=5) were administered with saline i.p. qd × 5, group C (n=5) were administered with 6-thioguanine 3 mg/kg i.p. qd × 5, group D (n=4) were administered with evofosfamide 50 mg/kg i.p. qd × 5/week for 3 weekly cycles. Tumours in group B-D were collected when the total tumour burden of one mouse in a group reached 4,000 mm³.

### Isolation of genomic DNA

For gRNA sequencing from cell cultures, 50 million cells were harvested from trypsinised cell suspensions, rinsed with PBS and stored at -80°C. Genomic DNA (gDNA) was isolated using the QIAamp DNA Blood Maxi Kit (QIAGEN). For tumour samples, harvested tumour tissue was cut into small fragments using a scalpel blade, and snap frozen in liquid nitrogen at 0.5 g/tube. Genomic DNA was isolated as described (S. Chen et al. 2015) with volumes scaled 3.5× used for up to 0.5 g of tumour tissue. Where tissue from a tumour exceeded 0.5 g, tissue was divided into tubes, gDNA was isolated from each tube and then pooled. Genomic DNA was desalted by adding NaCl to 0.2 M, followed by cold ethanol to precipitate the DNA and centrifugation at 18,000 × g to pellet the DNA. The pellet was rinsed once with cold freshly-prepared 70% ethanol, allowed to air-dry for 10 min and then dissolved in 10 mM Tris, pH 8.0 by heating to 50°C for 1 h. The gDNA was quality assessed by Nanodrop and agarose gel electrophoresis (0.8%/TBE), and quantified by Qubit DNA-BR assay (ThermoFisher Scientific).

### PCR

A three-stage PCR adapted from previous studies (S. Chen et al. 2015; Shalem et al. 2014) was used to amplify the gRNA sequences from the gRNA cassette in the gDNA, in preparation for Illumina sequencing. Herculase II Fusion DNA Polymerase (Agilent) was used for all reactions. Each reaction contained 0.5 µl DNA polymerase, 1 mM dNTP mixture, 4% DMSO and 1× Herculase II Buffer in 50 µl volume. Between each PCR, Ampure XP beads were used to purify the PCR products, and Qubit DNA-BR assay used to quantify products. In PCR1, 0.25 µM of forward and reverse primer (F1 and R2 from (Shalem et al. 2014)) and 5 µg gDNA per 50 µl reaction was used. Cycling parameters were an initial denaturation at 98°C for 5 min, 20 cycles of denaturation at 98°C for 60 s, annealing 62°C for 60 s and extension at 72°C for 90 s, and a final extension at 72°C for 10 min. For each cell culture sample at least 150 µg was used for PCR1 (split into individual reactions and pooled after PCR1) and for each tumour sample, at least 200 µg (or all gDNA available) was used. PCR2 was used to add the Illumina adaptors and barcode the samples, with 10 µl-equivalent of PCR1 mix and 0.2 µM of barcoded primers (PCR2 F Barcode 1-6 and R Barcode 1-6 from (S. Chen et al. 2015)) and 6 individual reactions performed for each sample. Cycling parameters were an initial denaturation at 98°C for 5 min, 10 cycles of denaturation at 98°C for 45 s, annealing 62°C for 35 s and extension at 72°C for 90 s, and a final extension at 72°C for 5 min. PCR3 was used to further amplify the product given low yields and amplification artefacts due to long primers in PCR2. PCR3 was performed with 0.2 µM of primers PCR3F (AATGATACGGCGACCACCGAGATC) and PCR3R (CAAGCAGAAGACGGCATACGAG), and 4 ng of purified PCR2 product. One reaction from pooled PCR2 product was performed. Cycling parameters were an initial denaturation at 98°C for 2 min, 10 cycles of denaturation at 98°C for 30 s, annealing 61°C for 35 s and extension at 72°C for 45 s, and a final extension at 72°C for 10 min. Primers were ordered from IDT, with HPLC-purified DNA oligos used for PCR1 and PCR3 and desalted Ultramer Oligos used for PCR2. Positive controls for PCR used lentiguide-Puro EGFP gRNA (Addgene 80036) which contained a gRNA sequence not present in the GeCKO library. Agarose gel electrophoresis (1.5%/TBE) was used to confirm PCR products (317 for PCR1, ∼360 for PCR2/PCR3). Samples were sequenced on a NextSeq500 (Illumina) using high-output, 2 × 150 bp flow cells (Illumina).

### Processing of sequencing data

A snakemake-based (Köster and Rahmann 2012) pipeline for processing sequencing data was developed (https://gitlab.com/twlee79/pooled_screen_counts). Sequencing data were demultiplexed according to reverse index by the Illumina platform and converted to FASTQ format. These data were then demultiplexed according to forward index using cutadapt v1.18 (Martin 2011) with non-internal adapter sequences, allowing up to 1 nucleotide mismatch. Fastqc 0.11.7 was carried out on the demultiplexed reads to check for sequencing quality. Next, cutadapt was used to trim poor quality bases and trim the vector 5’ and 3’ backbone sequence flanking the gRNA sequences. The quality threshold of 10 was used, and the 5’/3’ adapters were specified as linked adapters, allowing an adapter error rate of 0.1 and 19-21 nucleotide resulting sequences (expected 20-nucleotide gRNA sequences). The gRNA sequences were then aligned to the gRNA sequences present in the GeCKOv2 library using bowtie 2 (Langmead and Salzberg 2012; Langmead et al. 2019). Replicated gRNA sequences in the GeCKOv2 library were collapsed to individual entries prior to alignment (119,461 unique gRNAs).

Alignment scoring in bowtie 2 was set to local alignment mode with a score of 10 for a match, 4-6 for a mismatch (depending on quality) and per nucleotide gap penalty of -1, reporting the best alignment and also the best alternative alignment score. The number of reads aligned to each gRNA in the library were counted using a custom script (count_fgs_sam; https://gitlab.com/twlee79/count_fgs_sam). The minimum alignment score for a read to be counted was set to 189 – allowing for up to two mismatches or a 1-nt gap – with the additional criterion that only unambiguous alignments were counted. An unambiguous alignment was defined as a read where the main alignment had a score 3 or greater than the best alternative alignment, i.e. the main alignment had fewer mismatches or gaps than the best alternative; where the best alternative score differs by three or less, only the position of the mismatch or gap differ from the main alignment and this is considered an ambiguous alignment and excluded.

### Log counts per million

Given a dataset of gRNA read counts r_gi_ for samples *i* = 1, …, *N* and gRNAs *g* = 1, …, *G*, log counts per million (log-cpm) is defined as (Law et al. 2014):

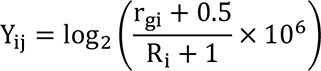

where R_i_ is the total number of counts for the sample (library size). The log-cpm is a measure of the proportion of counts in a sample that are for a given gRNA but is a poor measure of relative changes in the count for the gRNA across samples since the denominator (R_i_) is strongly affected by disproportionately large counts of certain gRNAs. To account for this, we calculated log normalised counts per million (log-ncpm), by replacing R_i_ with S_i_, the normalised library size for the given sample (see below).

### Read count normalisation

We devised a novel normalisation method, which we call MPM (mean of pairwise M values), to normalise our read count data by calculating normalised library sizes. This method is similar to the TMM (Robinson and Oshlack 2010) and RLE (Anders and Huber 2010) methods commonly used for RNAseq data, and GMPR (L. Chen et al. 2018) that was developed to deal with zero inflated counts in microbiome sequencing data. The general principle of these methods is that there are expected to be non-differential features (e.g. expressed genes or gRNAs) across samples. The non-differential features will be in the centre of the distribution of count ratios. Relative size factors between samples can be robustly estimated from the central non-differential features using a median or trimmed mean of count ratios.

In the MPM method, the influence of outlying high counts is reduced by clipping the top a% of counts in each sample – counts above the a-th percentile are replaced with the count at the a-th percentile. This ensures the normalisation is robust to large outlying counts and is analogous to ‘A’ trimming in the TMM method; trimming is replaced with clipping to retain all positive counts (in potentially zero-inflated data) for subsequent steps.

The pairwise log-fold-change (M) between samples i and j is calculated across gRNAs from clipped counts r’_gi_, with m% of valid ratios are discarded from each end to calculate the trimmed mean:

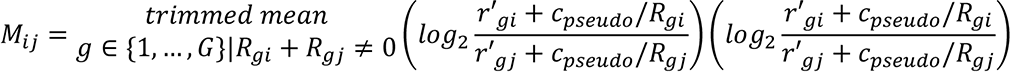

This is analogous to ‘M’ trimming in the TMM method. Note that only gRNAs that have zero counts in both samples are excluded from this calculation; gRNAs with a zero count in one sample only are used by offsetting the zero counts by the pseudo-count c_pseudo_, which we set as 0.5 as per the log-cpm definition above.

Although the use of a pseudo-count for zeroes has some weaknesses (L. Chen et al. 2018), their use here helps to better define ordering of count ratios prior to calculating the trimmed mean in spite of zero counts i.e. information that there was a lower count in one sample compared to the other is retained for the trimmed mean calculation. Furthermore, the influence of the pseudo-counts to the numeric value of M_ij_ is reduced by the use of the trimmed mean – since the pseudo-counts have a minimal value in one sample, a pseudo-count:actual count ratio tends to be more extreme and is more likely to be removed in the trim.

The size factor s_i_ for sample i on a log scale is calculated as the mean of M values to all other samples:

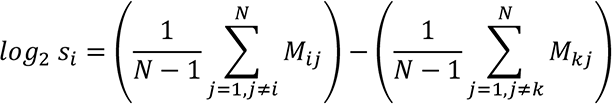

The second term scales the size factors to be relative to arbitrary sample k (i.e. s_k_ = 1). Note that unlike TMM, size factors are estimated from all pairwise ratios, as in RLE and GMPR. The size factor is a correction factor for library sizes, with normalised library size calculated as:

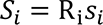

When normalised library sizes are used in place of (unnormalised) library sizes, log-ncpm becomes a measure of relative counts across samples. As the gRNA library contains 1000 non-targeting controls (NTCs), which can be assumed to be neutral, and thus non-differential on average, the size factors can be calculated using only NTC gRNAs i.e. *g* ∈ {*NTC gRNAs*} to give log-ncpm normalised to neutral gRNAs. In the study, we use MPM with a of 40%, m of 10% and NTC gRNAs only to calculate normalised library sizes.

### Binomial thinning simulations

To compare normalisation methods, we bootstrapped (sampled with replacement) the plasmid counts to a total of 1×10¹² reads. This bootstrapped data contained a median count 7.5×10⁶ per gRNA, necessary to allow individual gRNA counts of up to several million following binomial thinning. Next, we used binomial thinning using the seqgendiff package in R (Gerard 2020) to generate simulated datasets each containing 10 thinned columns (samples). In each dataset, a random signal was applied to expand/reduce counts of 95% of all gRNAs, generated using a normal distribution with mean 0 and standard deviation sampled from the uniform range 4.5-6.0; this signal is to account for stochastic clonal expansion/reduction. A fixed signal was also applied to all columns and to 100% of all targeting gRNAs. This signal was generated from a normal distribution with mean sampled from uniform [-1, -3] and standard deviation sampled from uniform [2, 3]. This signal is to account for guide-specific effects with a general dropout effect. Each column was further uniformly thinned to give a total number of reads uniformly sampled from the range [5e6, 15e6]. This uniform thinning causes many of the low counts to drop to zero and effectively zero-inflates the counts. These simulated datasets resembled the counts from our large tumour samples in terms of gRNA total counts, representation and bias. To ascertain the accuracy of each normalisation method tested, plasmid counts were appended to the simulated data and the normalisation method was executed. Next, the ratio of normalised counts per million of each simulated column compared to the plasmid column were determined for gRNAs with no signal (defined as combined random and fixed signal coefficients with absolute value of <0.01), and the median ratio was taken for each simulated column. This was repeated for a total of 32 random datasets with the median ratios pooled across the dataset and plotted to determine the normalisation method giving ratios closest to the expected ratio of 1 for these ’null’ gRNAs.

To determine statistical power, we performed binomial thinning on our small and large tumour datasets to simulate the effects of difference in counts between two equal-sized groups of samples using seqgendiff. To generate datasets containing fewer gRNAs, we randomly combined gRNAs into bins of 2, 5, 10, 20 and 50 gRNAs, with binning performed separately for targeting and non-targeting gRNAs and the counts of remaining gRNAs discarded to ensure all bins were derived from summing an equal number of gRNAs. We conducted analyses on either combined small tumours (total n=11 combining pilot UT-SCC-54C/NSG tumours and 14d tumours in treatment experiment) or large tumours (total n=28 combining no drug, 6-thioguanine and evofosfamide groups). The binned gRNAs were further subsampled to 100,000 (no binning), 50,000 (bin size = 2), 20,000, 10,000, 5000 and 2,000 (bin size = 50) gRNAs to ensure simulation outcome was not dependent on particular gRNAs. Samples were randomly divided into 2 groups, and binomial thinning was applied to add a log_2_ fold-change signal of N(0, 2²) to 20% of the targeting gRNAs/simulated gRNAs (bins). We then used MPM normalisation (to NTC counts) and voom/limma (Law et al. 2014) to determine which gRNAs could be detected as differentially reduced in on group versus the other at a false discovery rate (FDR) threshold of 0.10. For given effect sizes of applied signal, we determined the percentage of gRNAs with that effect size that were detected as being significantly different between the two groups as a measure of statistical power. We simulated 10 datasets per set of parameters. In each of these datasets, bin assignments, gRNA subsamples, group assignments and gRNAs receiving the signal were randomised.

## Results

### Selection of cell line for *in vivo* CRISPR/Cas9 screens

To establish an HNSCC tumour model suitable for CRISPR/Cas9 screening, we first sought to identify a HNSCC cell line with a high proportion of tumour-initiating cells. This was achieved by inoculating immunodeficient mice with different numbers of cells, to identify which of 23 HNSCC cell lines could initiate tumours with few cells. In initial experiments, only 11 of 23 UT-SCC cell lines grew as tumour xenografts within 100 days of inoculation of 5 × 10⁶ cells into NIH-III mice (Figure 1A). Four of these cell lines (UT-SCC-1B, UT-SCC-54B, UT-SCC-74A and UT-SCC-76A) had low tumour take rates and slow growth with ≤ 50% of tumours established by day 90, while two others (UT-SCC-1A and UT-SCC-42B) had frequent ulceration. There were no obvious correlations between tumour growth in mice and the primary site or lesion type of the HNSCC cell lines. Three of the seven UT-SCC cell lines that grew were from metastases, but there were two other lines from metastases that did not grow (Table 1).

**Figure 1.**
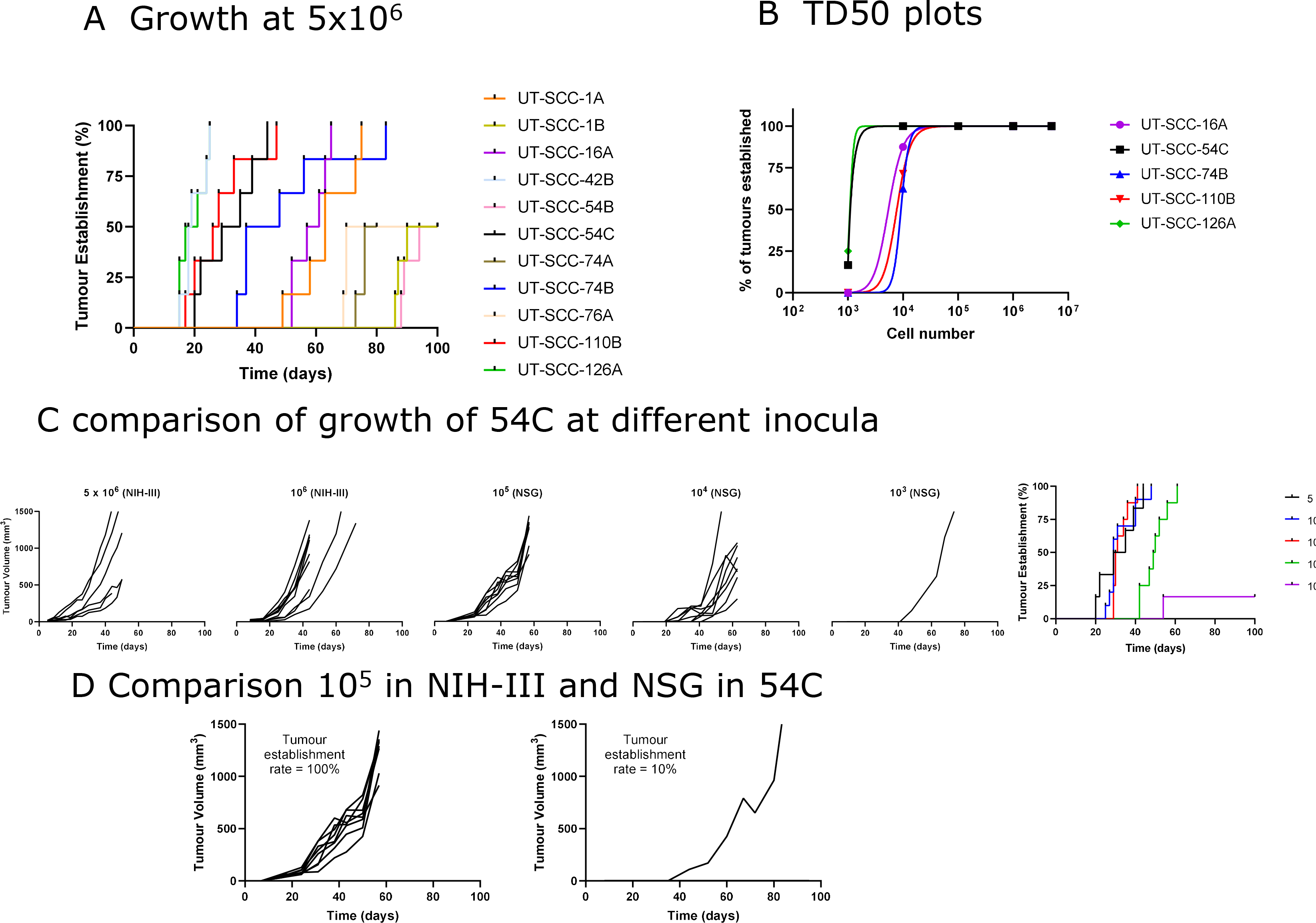
(A) Rate of tumour establishment for UT-SCC cell lines inoculated bilaterally in NIH-III mice at 5 × 10^6^ cells per flank. (B) Frequency of tumour establishment for UT-SCC cell lines inoculated into NIH-III or NSG mice at different cell inocula. (C) Comparison of UT-SCC-54C tumour growth in NIH-III and NSG mice at different cell inocula. The far right panel shows the rate of tumour establishment for UT-SCC-54C at different cell inocula. (D) Comparison of tumour establishment rate and growth for UT-SCC-54C tumours in NSG and NIH-III mice inoculated with 10^5^ cells.

**Table 1:** HNSCC cell line characteristics and TD_50_

| Cell Line | Primary Site | Type | TD50* | Ploidy** |
| --- | --- | --- | --- | --- |
| UT-SCC-54C | Buccal mucosa | Metastasis | 1100 | 2.00 |
| UT-SCC-126A | Labii inferioris | Primary | 1100; mice lose weight | 4.18 (53%), 3.70 (43%), 1.94 (4%) |
| UT-SCC-16A | Lingus | Primary | 5400 | 1.76 |
| UT-SCC-110B | Gingiva, maxillary sinus | Metastasis | 7700 | 3.07 |
| UT-SCC-74B | Lingus | Recurrent | 9200 | 2.72 (85%), 1.99 (15%) |
| UT-SCC-1A | Gingiva | Primary | <10<sup>6</sup>; ulcerates | 2.09 |
| UT-SCC-42B | Supraglottic larynx | Metastasis | <10<sup>6</sup>; ulcerates | 3.29 |
| UT-SCC-1B | Gingiva | Recurrent | Low take rate | 1.97 (92%), 3.13 (8%) |
| UT-SCC-54B | Buccal mucosa | Recurrent | Low take rate | 3.65 (98%), 1.89 (2%) |
| UT-SCC-74A | Lingus | Primary | Low take rate | 3.76 |
| UT-SCC-76A | Lingus | Primary | Low take rate | 3.18 |
| UT-SCC-16B | Lingus | Metastasis | Did not grow | 4.20 |
| UT-SCC-19A | Glottic larynx | Primary | Did not grow | 2.83 (70%), 3.76 (30%) |
| UT-SCC-19B | Glottic larynx | Recurrent | Did not grow | 2.36 |
| UT-SCC-24A | Lingus | Primary | Did not grow | 3.48 |
| UT-SCC-42A | Supraglottic larynx | Primary | Did not grow | 3.35 |
| UT-SCC-46A | Gingiva, maxilla | Primary | Did not grow | 2.08 |
| UT-SCC-54A | Buccal mucosa | Primary | Did not grow | 3.56 (97%), 1.76 (3%) |
| UT-SCC-59C | Parotid | Metastasis | Did not grow | Not determined |
| UT-SCC-60A | Tonsilla | Primary | Did not grow | **** |
| UT-SCC-63A | Gingiva, mandibula | Primary | Did not grow | 1.94 (50%), 3.56 (50%) |
| UT-SCC-76B | Lingus | Recurrent | Did not grow | 3.39 |
| UT-SCC-110A | Gingiva, maxillary sinus | Recurrent | Did not grow | 2.70 |

\* Low take rate indicates $\leq 50\%$ of tumours established by day 90 at $5 \times 10^6$ cells in NIH-III mice. Did not grow indicates no tumours were established by day 90 at $5 \times 10^6$ cells in NIH-III mice.
\*\* From Jamieson et al., 2018

The five cell lines with suitable growth kinetics (UT-SCC-1A, -16A, -42B, -54C, -74B, -110B, -126A) were selected to conduct limiting dilution assays in NIH-III and NSG mice to estimate the TD_50_, defined as the number of cells required for 50% tumour establishment, with a lower TD_50_ indicating a greater proportion of tumour-initiating cells. In these dilution experiments, tumour formation was assessed following inoculation at 10⁶ cells, and 10-fold reductions in cell number until fewer than 50% of tumours formed. Tumours were formed in 100% of NIH-III mice at ≥10^6^ for all cell lines, but the establishment rate fell with cell inocula ≤ 10^4^ in NSG mice (Figure 1B). As expected, tumour growth took longer on average with smaller inocula (Fig 1C) and tumour establishment rate was higher and faster in NSG mice than NIH-III mice (Fig 1D), and the lowest TD_50_ of approx. 1100 cells was estimated for UT-SCC-54C and UT-SCC-126A mice, followed by UT-SCC-16A (5400 cells), UT-SCC-110B (7700 cells) and UT-SCC-74B (9200 cells) (Table 1). Despite its low TD_50_ value UT-SCC-126A was associated with considerable weight loss in some animals (e.g. three of four NSG mice inoculated bilaterally with 10^4^ UT-SCC-126A required culling early due to ∼20% bodyweight loss).

Our earlier investigation had revealed that many of the cell lines in the UT-SCC panel were aneuploidy, with DNA content ranging from hypodiploid (DNA content of UT-SCC-16A = 1.76) to hypertetraploid (DNA content of UT-SCC-16B = 4.20) and some cell lines having subpopulations of cells with different DNA content (Jamieson et al. 2018) (Table 1). Our preference for our *in vivo* screen model was for cell lines that were pseudodiploid based on the reasoning that greater knockout efficiency would be achieved in such cell lines, with only two alleles on average needing knockout for recessive phenotypes versus four alleles on average for pseudotetraploid lines. Although previous results suggest gRNA potency is more important than target copy number in achieving efficient knockout, target copy number does have some effect (Yuen et al. 2017) and this remains a worthwhile criterion for selecting a cell line from a range of alternatives. Therefore, we choose to proceed with UT-SCC-54C cells, as they are pseudo-diploid (DNA content = 2.00) and have the joint lowest TD_50_, but without the weight loss concerns of UT-SCC-126A. For comparison, we selected UT-SCC-74B as a second model with a higher TD_50_ and that is predominantly hypotriploid (DNA content = 2.72 and 1.99). The in vitro doubling times of UT-SCC-54C and UT-SCC-74B were similar at 26.8 h and 29.7 h, respectively.

### Pilot xenograft experiments with UT-SCC-54C/Gecko and UT-SCC-74B/Gecko

Pilot *in vivo* screens with GeCKOv2 libraries were carried out in NSG and NIH-III mice to ascertain gRNA representation in small tumours and identify the optimal mouse strain. We first created Cas9-expressing pools of UT-SCC-54C and UT-SCC-74B cells, and transduced these at scale with the GeCKOv2 library.

Following selection for stable transductants using puromycin and expansion of the cell populations, we inoculated 10⁷ UT-SCC-54C/GeCKO and UT-SCC-74B/GeCKO cells into the flanks of NSG and NIH-III mice. Once the smallest tumour in each group reached ∼250 mm³, the tumours were harvested. UT-SCC-54C/GeCKO tumours grew faster in NSG male and female mice than NIH-III mice and faster than UT-SCC-74B/GeCKO tumours, which grew equally in NSG and NIH-III mice (Figure 2A).

**Figure 2.**
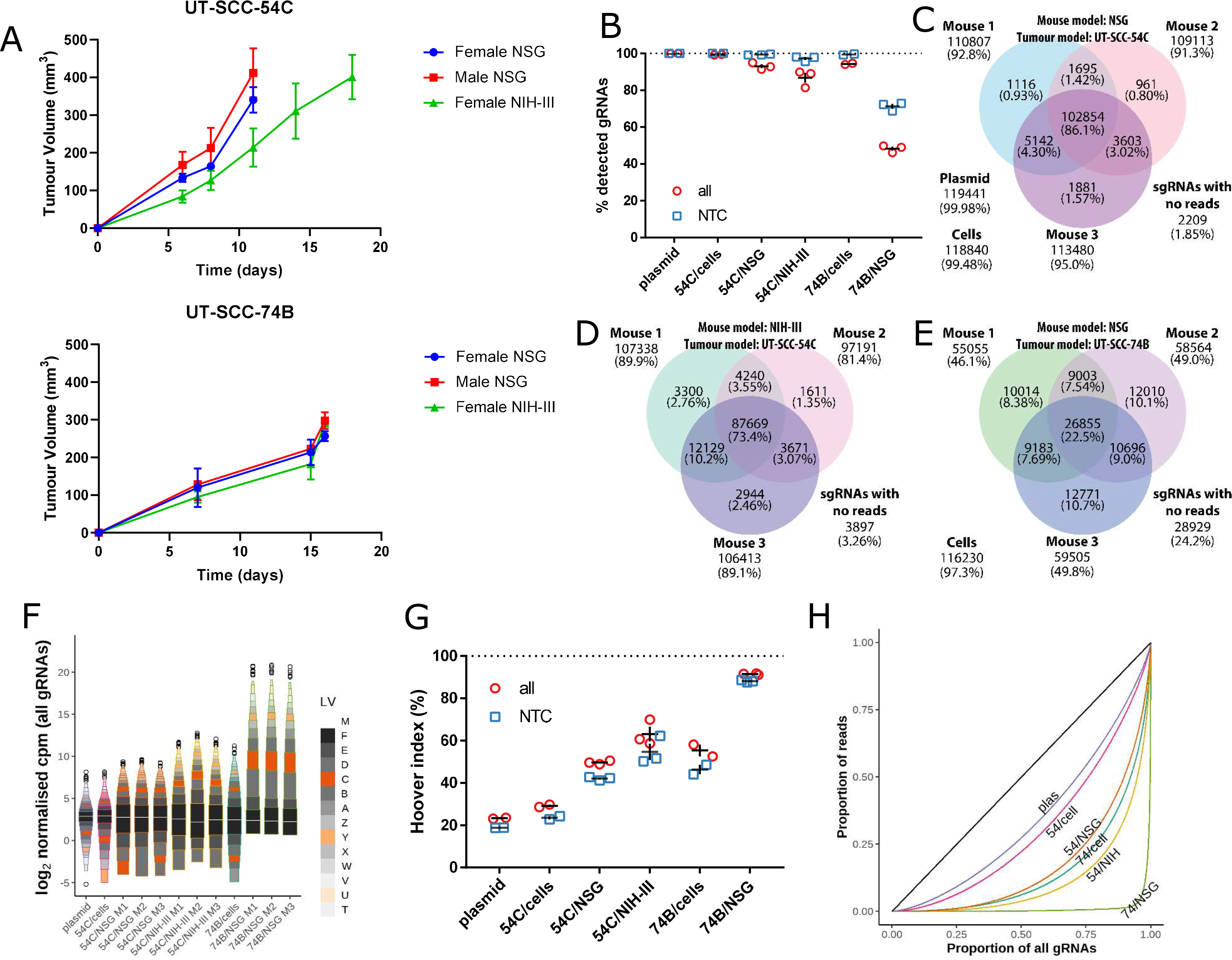
(A) Growth curves for UT-SCC-54C and UT-SCC-74B tumours in NSG and NIH-III mice. Bars represent the mean and SEM of 3-6 tumours. (B) Percentage of all or NTC gRNAs detected with counts of at least two for plasmid, UT-SCC-54C/GeCKO cell inoculum, UT-SCC-54C/GeCKO tumours in NSG mice, UT-SCC-54C/GeCKO tumours in NIH-III mice, and UT-SCC-74B/GeCKO cell inoculum and tumours in NSG mice. Replicates for plasmid and cell samples are PCR replicates from the same gDNA sample (n=2 each); tumour replicates are individual tumours in separate hosts (n=3 per group). The bar indicates mean ± SEM. (C-E) Venn diagrams showing overlap of detected gRNAs in UT-SCC-54C/GeCKO tumours in NSG mice, UT-SCC-54C/GeCKO tumours in NIH-III mice, and UT-SCC-74B/GeCKO in NSG mice. Values for plasmid and cell samples in (C) and (E) are for summed PCR replicates. (F) Letter-value plot (Hofmann, Wickham, and Kafadar 2017) of log normalised counts per million (log-ncpm) for all gRNAs in plasmid, cell inocula and GeCKO tumour samples. The white line inside the black boxes indicates the median (M), the black boxes indicate the upper/lower quartiles (F; fourths). The next smallest boxes indicate the upper/lower eighths (E) and so on. Counts for plasmid and cell samples were summed from two PCR replicates prior to normalisation. Remaining outliers are shown as open circles. (G) Hoover index of each sample as a measure of bias in the read counts for all and NTC gRNAs. (H) Lorenz curves to show distribution of all gRNA read counts. The curve for summed PCR replicates (plasmid and cell samples) or median tumour in a group is highlighted. The black line is the line of equality.

For determination of gRNA representation in the tumours, genomic DNA was isolated from tumours from female NSG and NIH-III mice, and the gRNA frequencies were read-out through a three-step PCR, Illumina sequencing and bioinformatics pipeline. Almost all gRNAs were detected at a threshold of 2 reads in the plasmid (99.98%) and UT-SCC-54C/GeCKO cells used for inoculation (99.5%; 2B, C). We observed high representation in UT-SCC-54C/GeCKO tumours in NSG hosts with 91.3-95.0% gRNAs detected (Figure 2B, C). This was somewhat lower in NIH-III hosts at 81.4-89.9% (Figure 2B, D). The vast majority of gRNAs were detected in UT-SCC-74B/GeCKO cells (97.3%) but UT-SCC-74B/GeCKO tumours in NSG mice had considerably lower representation of 46.1-49.8% (Figure 2B, E). In considering only non-targeting (NTC) gRNAs, >95% were detected in all UT-SCC-54C/GeCKO tumours and 69 to 73% in UT-SCC-74B/GeCKO tumours. This suggests that cells receiving neutral gRNAs had a growth or survival advantage within the tumours compared to cells receiving gRNAs targeting a gene (Figure 2B).

Our read count data, especially in UT-SCC-74B tumours, was zero-inflated at one end and contained very high counts for a few gRNAs at the other end.

Therefore, we devised a new normalisation procedure, which we call MPM, for our data to account for its unique characteristics (see methods). Following MPM normalisation, each sample had similar median normalised gRNA counts (Figure 2F), despite numerous zero counts and very high top counts (UT-SCC-74B/GeCKO tumours in particular). The more truncated spread of counts below the median is due to the floor of zero counts in the data (the lowermost boxes in Figure 2F are generally at zero counts). UT-SCC-74B/GeCKO tumours in NSG mice have size factors (ratio of normalised library sizes to the actual library size) of 0.016-0.019 (relative to the plasmid) and therefore much smaller normalised libraries sizes (approximately 200,000-300,000 compared to approximately 10 million for UT-SCC-54C NSG tumours) so the relative position of zero counts after normalisation is much higher (approximately 0 log-ncpm units) than for the other samples. This data representation reveals that the loss of representation from the plasmid to the cell libraries and the tumours is accompanied by a wider gRNA distribution. Among the tumour samples, the same pattern repeats, with the UT-SCC-54C/GeCKO tumours in NSG mice having higher representation and a narrower gRNA distribution than the UT-SCC-54C/GeCKO tumours in NIH-III mice, while the UT-SCC-74B/GeCKO tumours with the poorest representation had an extremely long upper tail indicating a highly biased gRNA distribution.

The same distribution pattern was also observed for the NTC gRNAs (Supp Fig 1A).

To further quantify the level of bias inherent in the gRNA distribution we determined the Hoover index for all gRNA and NTC gRNA counts (Figure 2G). The Hoover index can be interpreted as the percentage of reads that would have to be redistributed to other gRNAs to form a uniform distribution of gRNA counts. The Hoover index for all gRNA in UT-SCC-54C/GeCKO cells (28.6%) was slightly greater than that for the plasmid (23.1%). It increased to 48.9-50.5% for tumours in NSG mice and 58.6-70.0% for tumours in NIH-III mice. For UT-SCC-74B/GeCKO cells, the Hoover index was 54.0% increasing to 91.2-91.8% in NSG tumours. The NTC gRNA Hoover indexes followed the same pattern but were about 3-9% lower. The gRNA counts were also plotted as Lorenz curves (Figure 2H) to provide a more detailed graphical representation of bias in gRNA counts. The Lorenz curves for UT-SCC-74B/NSG, in particular, show the vast majority of counts dominated by the top 5% of gRNAs. These data suggest that the lower representation in the UT-SCC-74B/GeCKO tumours was driven by large differences in growth and/or survival in the cell population, with some cell clones expanding rapidly but others failing to do so. This growth/survival differential is unlikely to be primarily driven by gene knockouts as it was also observed to a similar but slightly lesser extent among cells containing neutral NTC gRNAs (Figure 2G; Supp Fig 1A-B).

### In vivo screens with UT-SCC-54C/Gecko

Based on these findings that UT-SCC-54C/GeCKO tumours had high gRNA representation without extreme levels of bias in gRNA count distribution, we reasoned that there may be an opportunity for selection for or against gRNA-induced knockouts to occur within these tumours. Therefore, we grew additional UT-SCC-54C/GeCKO tunours in female NSG mice. Tumours were allowed to grow until the median tumour size reached ∼250 mm³ then the mice were randomised into four groups: one (A) in which tumours were harvested immediately, a mock treatment group (B) and two drug treatment groups (C with 6-thioguanine, 6-TG, and D with evofosfamide, evo). Tumours in B-D were allowed to grow until the largest tumour in each group reached ethical size limits, at which point all tumours in the group were collected (day 38, 42 and 43 post inoculation for groups B, C and D, respectively). There was a small non-significant decrease in tumour growth for the two drug treatment groups compared to controls (Figure 3A). Tumour representation in the day 14 tumours was high (87.1 ± 1.4%; mean ± SEM), but was lower than in the similarly-sized tumours in the pilot study (93.0 ± 1.1%), most likely reflecting that the tumours grew slightly slower than in the pilot study, taking 14 days to reach an average size of 230 mm^3^ compared to 11 days to reach an average of 340 mm^3^.

**Figure 3.**
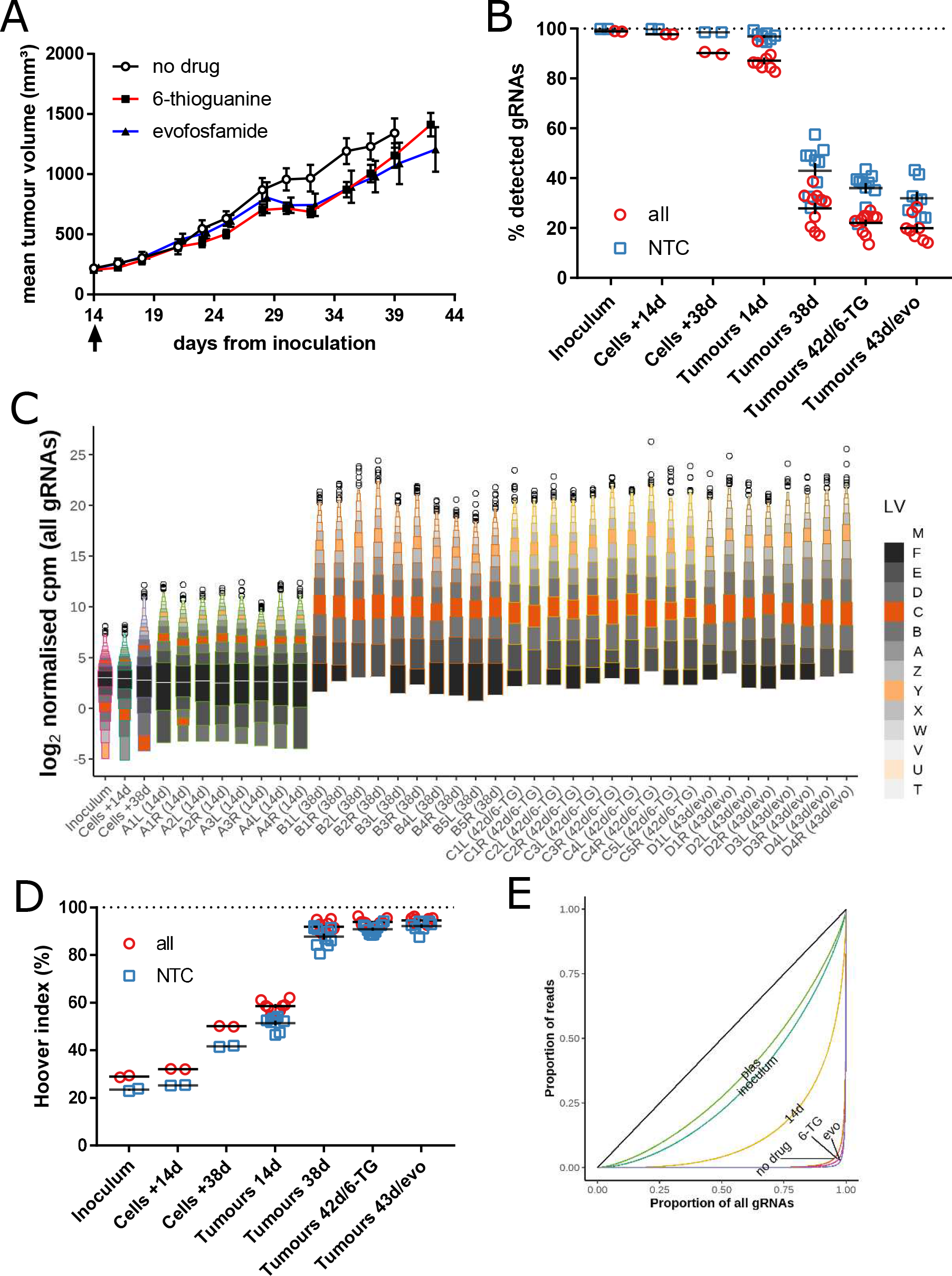
(A) UT-SCC-54C/GeCKO tumour growth after the commencement of drug treatment at day 14 (arrow) (mean ± SEM; n = 8-10). (B) Percentage of all or NTC gRNAs detected with counts of at least two for UT-SCC-54C/GeCKO cells (inoculum or cultured an additional 14 or 38 days after inoculation) and UT-SCC-54C/GeCKO tumours collected at 14 days, 38 days with no drug, 42 days with 6-thioguanine (6-TG) or 43 days with evofosfamide (evo). Replicate cell samples are PCR replicates from the same gDNA sample (n=2 each); tumour replicates are individual tumours grown bilaterally (n=8-10 per group). The bar indicates mean ± SEM. (C) Letter-value plot of log normalised counts per million (log-ncpm) for all gRNAs in cell and tumour samples. Counts for plasmid and cell samples were summed from two PCR replicates prior to normalisation. Sample name refers to group, animal number and flank position (e.g. A1L represents a tumour on the left flank of mouse 1 from group A) (D) Hoover index of each sample as a measure of bias in the read counts for all and NTC gRNAs. (E) Lorenz curves to show distribution of (all) gRNAs read counts. The curve for summed PCR replicates (plasmid and cell samples) or median tumour in a group is highlighted. The black line is the line of equality.

Moreover, there was a substantial decrease in representation in tumours in groups B, C and D to 27.9 ± 2.3%, 22.0 ± 1.4% and 20.0 ± 1.8% for all gRNAs (Figure 3B). NTC gRNAs decreased from 96.9 ± 0.6% at 14 days to 43.0 ± 3.2%, 36.0 ± 2.0% and 32.0 ± 2.5%, respectively. A much smaller loss of representation was observed by keeping the cells used for inoculation in culture with 97.7 ± 0.0 and 90.2 ± 0.5 of all gRNAs (and 99.9 ± 0.1 and 98.5 ± 0.0 NTCs) detected after 14 and 38 days in culture, respectively, compared to 98.9 ± 0.2 of all gRNAs (and 99.9 ± 0.0 NTCs) for the cell inoculum.

The distribution of normalised gRNA counts (Figure 3C; Supp Fig 2A for NTC gRNAs) showed a wider distribution of gRNA counts in 14-day tumours compared to the cell inoculum and cells cultured for an additional 14 days after inoculation but a similar distribution to cells cultured for an additional 38 days after inoculation. All large (38-43 day) tumours had gRNA distributions with very long upper tails indicating a strong bias towards relatively few gRNAs having very high counts while most others had zero or very few counts. The median gRNA count was zero for all large tumours. These biased distributions of gRNA counts are also indicated by the Hoover indices of 58.6 ± 0.8%, 92.0 ± 0.7%, 93.9 ± 0.4% and 94.6 ± 0.5%, for tumour groups A-D, respectively (Figure 3D). In comparison, the Hoover indices for the cell inoculum, +14 day and +38 day cultured cells were 28.7%, 31.8% and 50.0%, respectively. The Lorenz curves for these groups show the very high degree of bias in read counts that occurs in the large tumours, with 5-10% of all gRNAs accounting for >90% of all reads (Figure 3E).

### gRNA analysis of UT-SCC-54C/Gecko *in vivo* screen

The representation and bias data suggested a greater loss of targeting gRNAs compared to non-targeting controls. To more directly demonstrate this, we compared the ratio counts of all gRNAs to NTC gRNAs at the 90^th^ count percentile; the 90^th^ percentile was used to obtain a reasonable number of counts, since the median and upper quartile was zero for a number of large tumour samples. There was a decrease in the count ratio of all gRNAs compared to NTC gRNAs from the cell samples to the small tumours (14d), which dropped further in the large (38-43d) tumours (Figure 4A). To further investigate the performance of the MPM normalisation method on a more challenging dataset that was highly zero-inflated with very high read counts for certain gRNAs in the large tumour data, we performed simulations using binomial thinning to generate simulated datasets that resembled the large tumours. In these simulated datasets, we compared the actual normalised counts per million for a variety of normalisation methods to the expected value for gRNAs with no signal applied (expected ratio of 1). All normalisation methods were applied to counts of the NTC gRNAs alone, but corrected for total gRNA counts. We found that the total and upper quartile normalisation methods showed a strong negative bias (Figure 4B) due to the inflated counts of certain gRNAs. The TMM (Robinson and Oshlack 2010) and RLE (Anders and Huber 2010) methods commonly used for RNAseq also showed a slight negative bias, with the RLE method performing better than TMM. In these methods, any zero counts are excluded from the data. Setting zeroes to a pseudo-count of 0.5 reduced the bias with the TMM method and virtually eliminated it for the RLE method. The GMPR method, which is designed for zero-inflated data from microbiome sequencing projects (L. Chen et al. 2018) showed only a slight negative bias. Our MPM method showed very little bias among these simulated data and was generally robust to different parameters, although is computationally more expensive than the similarly performing RLE/pseudo method due to the calculation of all pairwise ratios.

**Figure 4.**
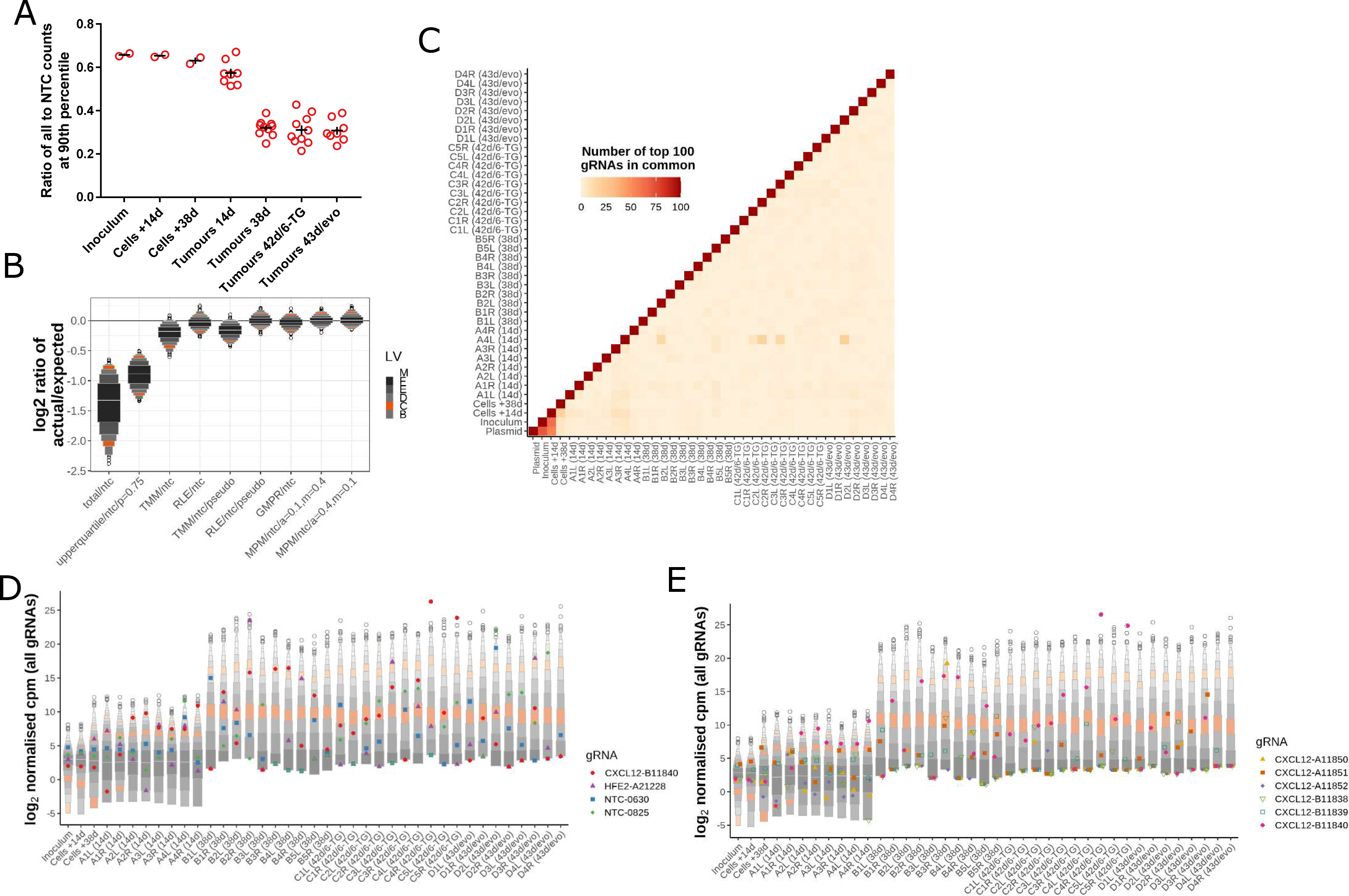
(A) Ratio of counts for the gRNA at the 90th count percentile among all gRNAs to counts for the NTC gRNA at the 90th count percentile. The bar indicates mean ± SEM. (B) Letter-value plots of log_2_ ratio of actual normalised counts per million to expected counts per million for different normalisation methods. Data are derived from binomial thinning simulations of datasets with characteristics of large tumours. ‘Total’ is normalisation to the total number of NTC gRNAs, ‘upperquartil’ is normalisation to the count at the upper quartile of NTC gRNAs. The ‘pseudo’ versions of the TMM and RLE methods were performed after setting a pseudo-count of 0.5 to all zero counts. All normalisations were applied to the NTC gRNA counts alone. The letter-value plots summarise the actual/expected ratio of the median null gRNA for each simulated sample. The expected ratio is 1 (0 on a log scale), so the best methods have a distribution closest to 0. (C) Heat map showing the number of the top 100 gRNAs with the highest counts that are shared among each pair of samples. The high number (red) indicates that the top gRNAs in each pair of samples are similar, while a low number (white) indicates each pair of samples has different top gRNAs. (D) Counts of four gRNAs across samples shown on a letter-value plot of log normalised counts per million (log-ncpm) for all gRNAs in cell and tumour samples. The four gRNAs represent those with the highest and 10th highest maximum log-ncpm across all samples for all or NTC gRNAs. (E) Counts of the six gRNAs in the library targeting CXCL2 shown on a letter-value plot of log-ncpm for all gRNAs in cell and tumour samples.

We further analysed whether the gRNAs with the most reads in the tumour samples tended to be the same across samples within a group, indicative of gene knockout altering tumour growth or drug therapy. However, instead, we found that the top gRNAs were different in each sample (Figure 4D). This indicates that the highly expanded counts among these top gRNAs was primarily driven by gRNA-driven selection, but rather random expansion of certain clones within the tumours. To further explore this, we investigated the read counts of the gRNAs with the highest (CXCL12 gRNA #B11840) and 10^th^ highest (HFE2 gRNA #A21228) log-ncpm across all samples, as well as the NTC gRNAs with the highest (NTC gRNA #0825) and 10^th^ highest (NTC #0630) log-ncpm across all samples. The read counts of these four gRNAs varied widely across large tumour samples (Figure 5D). For instance, the log-ncpm for CXCL12 gRNA #B11840 in one 6-TG-treated tumour was 26.5 while zero counts were detected in another 6-TG-treated tumour. This wide variation suggests largely random expansion of particular clones within these tumours. In smaller tumours the counts were more consistent but were still unusually high or low in certain samples, indicating a degree of random clonal expansion was already present in these smaller tumours. To confirm this variation applied across all gRNA targeting the same gene, we also investigated all gRNA targeting CXCL12 and observed a similar level of variation between samples as we had seen for CXCL12 gRNA #B11840 (Figure 4E).

**Figure 5.**
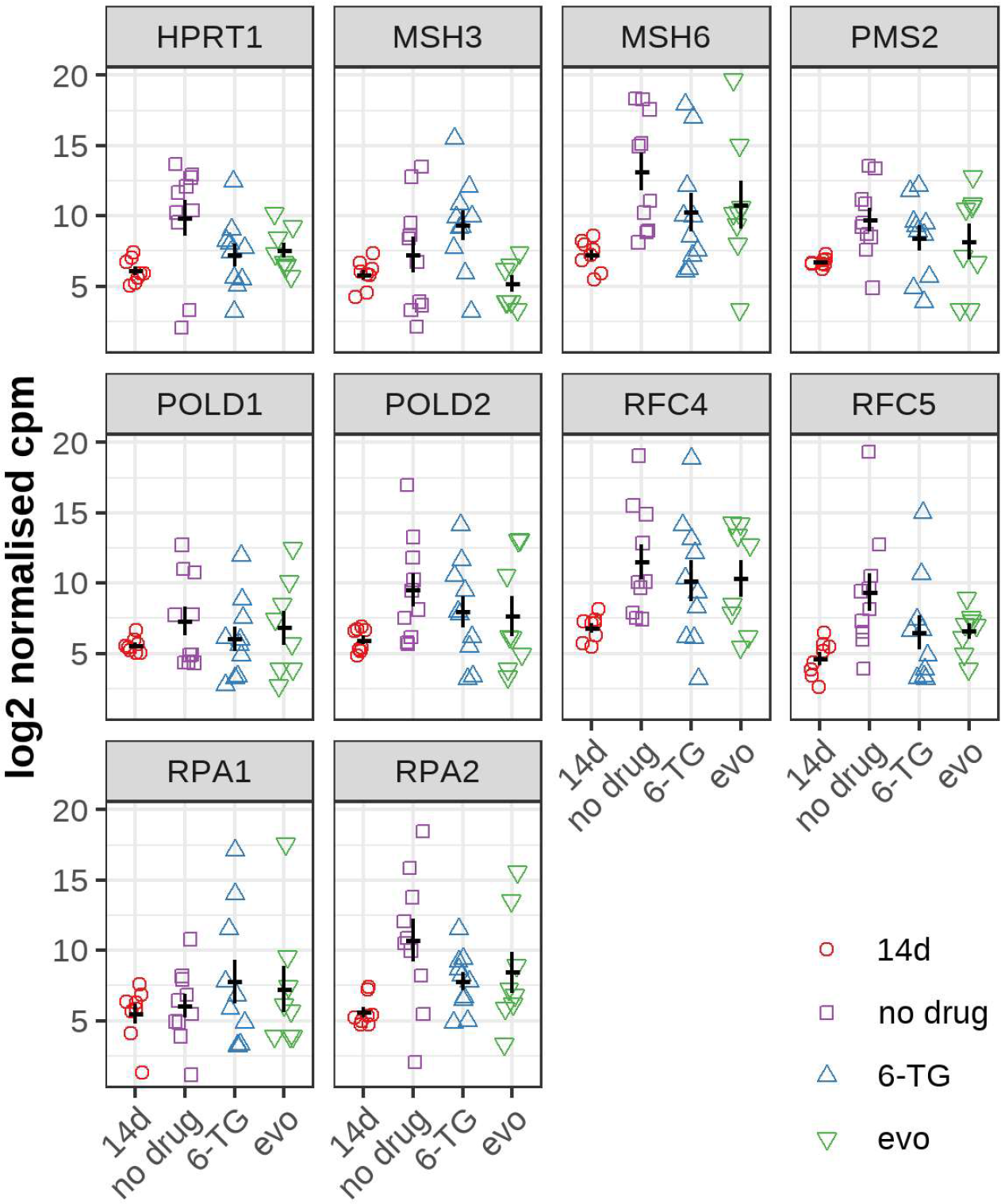
Log normalised counts per million (log-ncpm) for the gRNAs targeting genes expected to be involved in 6-thioguanine resistance in the tumour samples. The counts of all gRNAs targeting these genes were summed prior to normalising. The selected genes were *HPRT1, NUDT5* and genes in the KEGG mismatch repair geneset. Counts for ten genes with the highest log fold-change difference between no drug and 6-TG shown. The bar indicates mean ± SEM.

The large degree of random variability in read counts of individual gRNAs in the large tumours, precluded the applicability of conducting gene level analyses to identify genes that influence tumour growth or response to therapy. We confirmed this by analysing the 6-TG-treated tumours to determine if the well-characterised 6-TG sensitivity genes HPRT1 and NUDT5 (Doench et al. 2016), as well as those in the mismatch repair pathway (T. Wang et al. 2014), were enriched in treated vs control tumours. Given the large variation in individual gRNA counts highlighted previously (Figure 4D-E), we summed the counts for all gRNAs targeting each of these genes prior to plotting the log-ncpm of these summed gRNA counts (Figure 5). Despite a reduction in variation through the use of summed counts, the log-ncpm within each group varied widely, and was greater than the variation between groups. Due to the large within-group variance, it was not possible to determine the enriched or depleted gRNA clones between the untreated and treated groups with statistical confidence.

### Simulation of suitable library size for *in vivo* screen

The GeCKOv2 whole genome gRNA library used in this study consists of 119,461 unique gRNAs. Therefore, the random clonal expansion would need to include >100,000 clones for this process to not bias gRNA representation. A smaller gRNA library in tumours in theory would allow greater averaging per gRNA and increase gRNA representation in random clonal expansion. To assess whether a smaller gRNA library would be more suitable for gene level analyses in *in vivo* screens, we simulated smaller gRNA libraries by randomly combining the gRNA counts in our full-size library into bins of 2, 5, 10, 25, and 50 gRNAs. We reasoned that our binned counts would mimic the random characteristics of clonal expansion in smaller libraries, as clones with different gRNAs in the larger library would be largely equivalent (in terms of random characteristics) to different clones that receive the same gRNA in a smaller library. Any signal due to the gRNAs themselves, which we have already established was small relative to the random clonal expansion effect (Figure 5), was further averaged out by the random nature of the binning, particular among bins with a greater number of gRNAs. Binning into groups of 50 gRNAs, equivalent to about 2400 gRNAs, tightened the range in gRNA log-ncpm among large tumour samples to a median of 15.2 (range 11.7-21.5; Figure 6A), similar to small tumours for unbinned data (median 15.5; range 14.1-16.3). The representation was 100%, and the median Hoover indexes was 63.3% (range 49.7-73.0%), again similar to that of small tumours (Figure 3D). To assess the variation in counts of individual gRNA bins, we plotted the log-ncpm of the gRNA bins with the highest and 10^th^ highest log-ncpm across all samples for targeting and non-targeting gRNAs (Figure 6A). The variation in log-ncpm for these gRNA bins was considerably reduced compared to unbinned gRNAs in large tumours (Figure 6D-E) and was similar in extent to that in small tumours. This demonstrates that averaging over a greater number of clones per gRNA, through the use of a smaller gRNA library, could partly counteract the variance expansion effect of stochastic clonal expansion that occurs in the tumours.

**Figure 6.**
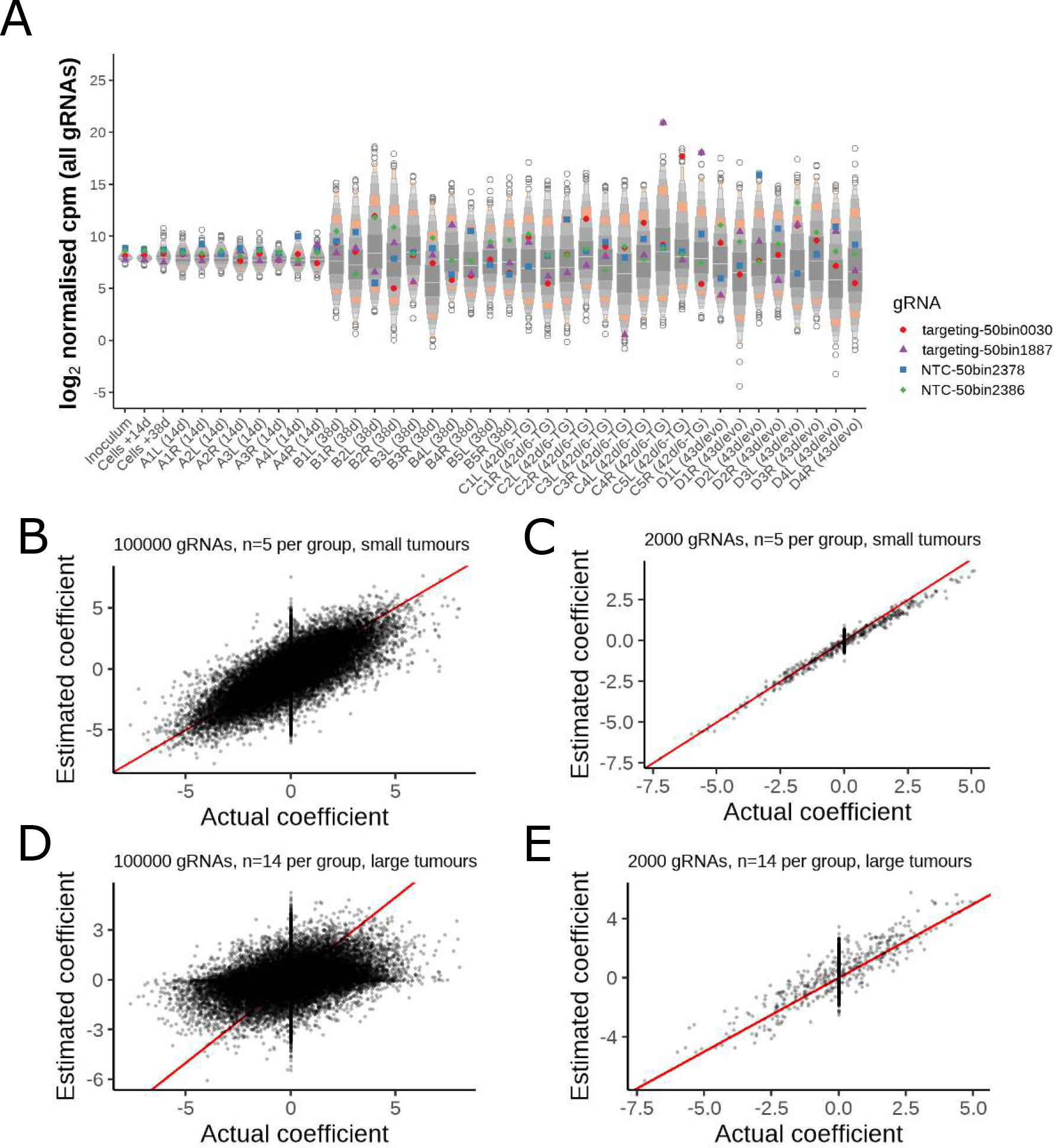
(A) Counts of four gRNA bins across samples shown on a letter-value plot of log normalised counts per million (log-ncpm) following random binning of gRNAs into bins of 50 in cell and tumour samples. The four gRNA bins represent those with the highest and 10th highest maximum log-ncpm across all samples for all or NTC gRNAs and are named arbitrarily. (B) Estimated coefficients (log_2_ fold-change between groups) from voom-limma analysis in a two-group dataset simulated by binomial thinning by applying a difference between the two groups. The estimated coefficient against the actual applied coefficient is shown, with the red diagonal line indicating equality between the two. The signal was applied to 20% of targeting gRNAs only, so the spread of points at the actual coefficient of zero represents the variation of estimated coefficients for null gRNAs. The dataset was simulated by subsampling the unbinned gRNA data for small tumours to 100,000 gRNAs, randomly assigning samples to groups of n=5 and applying binomial thinning using N(0,2²). (C) Estimated versus actual coefficients for a simulated dataset from small tumours with gRNAs binned into groups of 50 and subsampled to 2,000 gRNA bins and n=5 samples randomly assigned per group. (D) Estimated versus actual coefficients for a simulated dataset from large tumours with unbinned gRNAs subsampled to 100,000 gRNAs bins and n=14 samples randomly assigned per group. (E) Estimated versus actual coefficients for a simulated dataset from large tumours with gRNAs binned into groups of 50 and subsampled to 2,000 gRNA bins and n=14 samples randomly assigned per group.

To assess whether the use of a smaller gRNA library would facilitate the detection of differences in gRNA abundance between two groups, we carried out simulations using binomial thinning (Gerard 2020). Simulation using binomial thinning ensures that the characteristics of the original dataset are retained, in contrast to the use of theoretical models, and therefore provides a more realistic assessment of ability to detect differential changes in gRNA abundance in these data. Note that the signal applied in the binomial thinning method is done by decreasing counts in one group while the counts in the other group are kept constant, thus it represents a dropout signal or negative selection (binomial thinning does not expand counts and therefore cannot simulate positive selection). In our simulations, we tested the ability to detect an applied signal using voom/limma (Law et al. 2014) on our gRNA datasets from small tumours and large tumours, while varying the group size (number of tumours per group) and number of gRNAs (through binning and subsampling). In simulations on the unbinned small tumour dataset, voom/limma analysis was able to estimate the applied signal to a reasonable degree (Figure 6B). However, this estimated signal fell within the range of noise in the dataset, as indicated by the spread of estimated coefficients for null gRNAs (actual coefficient = 0), which would make detecting statistically significant differences challenging except at the extremes of the applied signal (e.g. ±5). Following binning/subsampling to a dataset of 2,000 simulated gRNAs, the signal was more precisely estimated and there was also a large decrease in the variance of null gRNAs (Figure 6C). We determined the statistical power from these simulations when varying the number of gRNAs in the dataset (Table 2), which showed that differences of 1-2 log_2_ fold-change between groups could be detected with 90% power in datasets of 2,000 gRNAs and n=5 tumours per group at FDR<0.1. Statistical power was also high with up to 10,000 gRNAs and moderate with 20,000 gRNAs, but low for 50,000 or 100,000 gRNAs. The false discovery proportion (at null signal) was well below 0.1 indicating conservative control of FDR by voom/limma.

**Table 2.** Statistical power determined by binomial thinning simulation in the small tumour dataset at a group size of n=5 tumours/group, varying the number of gRNAs by random binning/subsampling and for different levels of applied signal. The median percentage of gRNAs across simulations (range in brackets) detected at a false discovery rate of 0.1 or below for each level of applied signal is provided as an estimate of statistical power. For null gRNAs, this is the false discovery percentage.

| Dataset: Group size n=5, small tumours |  |  |  |  |  |  |
| --- | --- | --- | --- | --- | --- | --- |
| % of gRNAs detected with FDR<0.1 at each effect size (log2 FC) |  |  |  |  |  |  |
| Number of gRNAs | null (<0.01) | [0.5-1) | [1-2) | [2-3) | [3-4) | ≥4 |
| 2000 | 1.3 (0.7-1.8) | 47.3 (44.7-48.7) | 91.1 (86.4-93.9) | 100 (100-100) | 100 (100-100) | 100 (100-100) |
| 4000 | 1.1 (0.9-1.2) | 22.1 (19.7-26.0) | 73.2 (68.8-80.4) | 98.4 (97.7-99.3) | 100 (100-100) | 100 (100-100) |
| 10000 | 0.7 (0.6-0.8) | 9.5 (8.5-10.7) | 43.0 (41.6-45.5) | 83.5 (82.5-84.6) | 97.9 (96.0-99.4) | 100 (100-100) |
| 20000 | 0.4 (0.4-0.4) | 3.2 (2.2-4.4) | 17.3 (15.1-19.9) | 53.9 (51.0-57.3) | 84.5 (84.1-84.9) | 95.6 (93.6-96.9) |
| 50000 | 0.0 (0.0-0.1) | 0.1 (0.1-0.1) | 0.9 (0.2-1.4) | 5.4 (2.3-7.5) | 18.3 (8.9-24.4) | 42.1 (28.3-50.3) |
| 100000 | 0.1 (0.0-0.1) | 0.1 (0.1-0.2) | 0.4 (0.3-0.5) | 1.6 (1.3-1.8) | 7.1 (6.4-8.1) | 24.9 (21.8-27.0) |

In simulations in the dataset of large tumours, with 100,000 gRNAs and n=14 per group, the applied signal was not able to be estimated (Figure 6D) and there was a very large degree of noise present among null gRNAs.

Binning/subsampling to 2,000 simulated gRNAs allowed the applied signal to be estimated reasonably precisely, although there was still quite a high degree in noise among the null gRNAs (Figure 6E). With a group size of n=7, statistical power was estimated as >30% with effect sizes of 3-4 or greater with 2,000 gRNAs, but low (<10%) with smaller effect sizes or in datasets containing >10,000 gRNAs (Table 3). With 20,000 gRNAs or greater, even the strongest applied signals (effect size ≥4) could not be detected in the simulations. When group size was increased to n=14 in large tumours, statistical power was moderate at 47% to detect effect sizes of 2-3 log_2_ fold-change at FDR<0.1 in datasets of 2,000 gRNAs, with high power (>80%) to detect larger effects.

**Table 3.** Statistical power determined by binomial thinning simulation in the large tumour dataset at a group size of n=7 tumours/group, varying the number of gRNAs by random binning/subsampling and for different levels of applied signal. Statistical power as median percentage of gRNAs (range in brackets) detected at FDR of <0.1 is shown.

| Dataset: Group size n=7, large tumours |  |  |  |  |  |  |
| --- | --- | --- | --- | --- | --- | --- |
| % of gRNAs detected with FDR<0.1 at each effect size (log2 FC) |  |  |  |  |  |  |
| Number of gRNAs | null (<0.01) | [0.5-1) | [1-2) | [2-3) | [3-4) | ≥4 |
| 2000 | 0.1 (0.0-0.2) | 0.8 (0.0-1.2) | 1.4 (0.8-1.7) | 9.0 (4.5-15.7) | 31.2 (25.8-35.3) | 49.8 (46.2-53.3) |
| 4000 | 0.1 (0.0-0.2) | 0.2 (0.0-0.6) | 0.8 (0.0-2.0) | 4.5 (0.8-8.0) | 14.4 (4.8-23.1) | 37.6 (25.7-50.0) |
| 10000 | 0.0 (0.0-0.1) | 0.2 (0.0-0.3) | 0.2 (0.2-0.3) | 1.2 (0.3-2.4) | 3.9 (0.0-7.2) | 14.0 (10.7-18.6) |
| 20000 | 0.0 (0.0-0.0) | 0.0 (0.0-0.0) | 0.0 (0.0-0.0) | 0.0 (0.0-0.0) | 0.0 (0.0-0.0) | 0.0 (0.0-0.0) |
| 50000 | 0.0 (0.0-0.0) | 0.0 (0.0-0.0) | 0.0 (0.0-0.0) | 0.0 (0.0-0.0) | 0.0 (0.0-0.0) | 0.0 (0.0-0.0) |
| 100000 | 0.0 (0.0-0.0) | 0.0 (0.0-0.0) | 0.0 (0.0-0.0) | 0.0 (0.0-0.0) | 0.0 (0.0-0.0) | 0.0 (0.0-0.0) |

**Table 4.** Statistical power determined by binomial thinning simulation in the large tumour dataset at a group size of n=14 tumours/group, varying the number of gRNAs by random binning/subsampling and for different levels of applied signal. Statistical power as median percentage of gRNAs (range in brackets) detected at FDR of <0.1 is shown.

| Dataset: Group size n=14, large tumours |  |  |  |  |  |  |
| --- | --- | --- | --- | --- | --- | --- |
| % of gRNAs detected with FDR<0.1 at each effect size (log2 FC) |  |  |  |  |  |  |
| Number of gRNAs | null (<0.01) | [0.5-1) | [1-2) | [2-3) | [3-4) | ≥4 |
| 2000 | 0.7 (0.6-0.8) | 1.3 (1.2-1.5) | 13.4 (12.0-15.7) | 47.2 (44.1-50.0) | 84.7 (75.0-100) | 94.8 (88.9-100) |
| 4000 | 0.3 (0.3-0.3) | 1.1 (0.7-1.4) | 8.3 (7.3-10.0) | 30.0 (27.2-33.6) | 64.3 (59.7-67.6) | 89.5 (77.4-97.2) |
| 10000 | 0.2 (0.1-0.3) | 0.2 (0.0-0.3) | 2.8 (2.2-3.4) | 12.5 (10.2-13.7) | 30.9 (28.6-33.5) | 58.7 (53.5-61.3) |
| 20000 | 0.0 (0.0-0.1) | 0.1 (0.0-0.4) | 0.6 (0.2-0.9) | 2.0 (1.3-2.9) | 7.9 (6.6-10.1) | 20.5 (17.2-24.3) |
| 50000 | 0.0 (0.0-0.0) | 0.0 (0.0-0.0) | 0.0 (0.0-0.0) | 0.0 (0.0-0.0) | 0.0 (0.0-0.0) | 0.0 (0.0-0.0) |
| 100000 | 0.0 (0.0-0.0) | 0.0 (0.0-0.0) | 0.0 (0.0-0.0) | 0.0 (0.0-0.0) | 0.0 (0.0-0.0) | 0.0 (0.0-0.0) |

Power dropped to 30% with 4,000 gRNAs and 12.5% with 10,000 gRNAs. With 20,000 gRNAs or greater, statistical power was ≤2% . Again, the false discovery proportion indicated conservative control of FDR. Altogether, these simulations indicate that changes in gRNA abundance due to selection effects would be detectable if the degree of random clonal expansion is controlled either by the use of small tumours (or those grown for a short period of time), and/or the size of the gRNA library is reduced to 4,000 gRNAs or fewer to ensure greater averaging of clones in the library to counteract the large count variance that occurs due to random clonal expansion.

## Discussion

In vivo CRISPR/Cas9 screens offer the opportunity to identify genes responsible for a specific phenotype in response to a selective pressure, while maintaining the complex multicellular interactions of the tumour microenvironment. In this study, we sought to establish a tumour xenograft model with high gRNA representation suitable for whole genome *in vivo* CRISPR/Cas9 screens. To do this, we first inoculated immunodeficient mice with different cell numbers of multiple early-passage human HNSCC cell lines to identify a tumour xenograft model with a low TD_50_ indicating a high frequency of tumour-initiating cells. We then selected low and moderate TD_50_ lines for transduction with the GeCKOv2 whole-genome gRNA library and compared gRNA representation in small tumours generated in NSG mice by inoculation of the transduced cells. Our low TD_50_ UT-SCC-54C line retained >99% gRNA representation in cultured cells, presumably owing to its diploid nature, and had much higher gRNA representation than the moderate TD_50_ cell line UT-SCC-74B when GeCKO-transduced cells were inoculated into NSG mice at 10^7^ cells. Although not-definitive since only two models were used, our data does suggest that using a conducive mouse host and cell line with a high frequency of tumour-initiating cells can lead to high gRNA representation on tumour initiation.

Despite the high gRNA representation at tumour initiation with our UT-SCC-54/GeCKO tumour model, a considerable loss in gRNA representation was observed as tumours were grown to >1 cm^3^, reflecting similar outcomes of previous whole-genome *in vivo* screens (S. Chen et al. 2015; Song et al. 2017; Rushworth et al. 2020; Kodama et al. 2017). By evaluating the gRNA with the highest read counts in the large tumours as well as the read counts of known 6-TG sensitivity genes in the 6-TG-treated tumours, it was apparent that the loss of representation occurred due to stochastic clonal expansion. Our study was not set up to evaluate Darwinian clonal selection, since pre-existing clonal heterogeneity independent of gRNA-mediated gene knockout may have been present in the UT-SCC-54C/GeCKO cells. Yet, if clonal selection was the major evolutionary driver, we would have expected greater reproducibility in gRNA read counts between tumour samples. Indeed, previous work has highlighted that for the majority of xenograft models, a small number of clones rapidly expand to eventually dominate the tumour as it grows (Ben-David et al. 2017; Eirew et al. 2015). For our model at least, this clonal expansion expedited by the genetic bottleneck of tumour initiation (Ben-David et al. 2017) is dependent more on the tumour-initiating ability of the cell, rather than gRNA-enforced gene knockout.

The random clonal expansion that occurred with tumour progression in our xenograft model greatly increased the variance in gRNA counts and was the major limiting factor in identification of drug sensitivity genes as any gRNA-driven effects would require very high effect sizes to be detected. This highlights the major technical challenges associated with using whole-genome gRNA libraries in transplantable *in vivo* screens. We utilised the human GeCKOv2 full gRNA library of 119,461 unique gRNAs in our study with six gRNA on average against each gene. Although other groups have also used the human or mouse GeCKOv2 library for whole-genome screens, in each case the half-library of ∼65,000 gRNA was used with only three gRNA on average against each gene(S. Chen et al. 2015; Song et al. 2017; Rushworth et al. 2020; Yau et al. 2017; Dai et al. 2021; Kodama et al. 2017). Interestingly, the two of these studies that reported the highest detection of gRNA in tumours inoculated 3 × 10^7^ cells into mice (Yau et al. 2017; Dai et al. 2021), a 3-fold increase on our inoculum. This extends to a ∼6-fold increase in cells/gRNA (∼450 vs ∼83 in our study) when the half-library is factored in and may have contributed to greater averaging across clones in tumours and a reduction in count variance compared to our study. The use of only three gRNA on average reduces the statistical confidence that any detected change is a true discovery, but nevertheless is likely a necessary compromise to increase the chance of a successful whole-genome screen. A further factor that may have contributed to the success of these studies compared to ours is the particular growth characteristics of the cell line in vivo. The degree of gRNA loss and bias in small (14d/250 mm³) UT-SCC-74B/GeCKO tumours in our study was similar to that of large (∼40d/>1000 mm³) UT-SCC-54C/GeCKO tumours, with the use of NSG over NIH-III hosts providing only a minor improvement. It is possible the particular cell line/host combinations provide more suitable tumour growth characteristics (i.e. more tumour initiating cells) for *in vivo* screens than others, although this likely needs to be empirically determined by pilot xenograft experiments of the cell library carried through to likely tumour collection times/sizes following drug treatment.

Our findings suggest other possible improvements that can be implemented to increase the ability of whole-genome *in vivo* CRISPR/Cas9 screens to uncover drug sensitivity genes. Early tumours were found to have greatly reduced gRNA loss and stochastic expansion compared to late tumours. This suggests that initiating tumour treatment when tumours are small (∼100 mm^3^) and collecting tumours 7-14 days after treatment when tumours at most are moderate in size (<750 mm^3^) may help to avoid the dramatic stochastic expansion that occurs with big tumours that have been left to grow for several weeks. Our simulation studies indicated a large difference in statistical power between small and large tumours due to cumulative effects of stochastic expansion in the large tumours, and that considerable gains in statistical power can be achieved by minimising time/growth-dependent stochastic effects. The quality of the gRNA library will also affect its ability to uncover sensitivity genes. The GeCKOv2 library has been demonstrated to have reduced performance than newer gRNA libraries, due to a larger number of poorly functioning gRNA (Doench et al. 2016; Sanson et al. 2018). Switching to an optimised whole-genome library, e.g. Brunello and Brie libraries, that have fewer gRNA on average per gene than GeCKOv2, but all gRNA efficiently target their intended gene with minimal off-target activity (Doench et al. 2016; Sanson et al. 2018) will ensure the effect sizes of gRNA knockout are not hampered by inefficient knockout, increasing power to detect drug sensitivity genes. Another potential approach to improve representation of whole-genome *in vivo* CRISPR/Cas9 screens is to transduce multiple gRNA per cell (Weber et al. 2020), especially if each gRNA had a unique barcode (Zhu et al. 2019) to differentiate their phenotypic effects.

The optimal approach to ensure success in *in vivo* CRISPR/Cas9 screens, however, is to reduce library size. Our binomial thinning simulations revealed that although our UT-SCC-54C/GeCKO xenograft model was not suited for identifying drug sensitivity genes with a >100,000 gRNA library, if the gRNA library was reduced to ∼2,000 gRNA, the effects of stochastic expansion were greatly reduced in the model so that there is sufficient statistical power (∼40%) to identify some drug sensitivity genes provided group sizes are large (approx. n=15). Our simulations also showed that the use of small libraries (2,000-4,000 gRNAs) also greatly improves statistical power in small tumours, so that there is high power to detect drug sensitivity genes with a modest group size (n=5).

Owing to their improved data quality, smaller libraries are the method of choice for most *in vivo* CRISPR/Cas9 screens (Manguso et al. 2017; Li et al. 2020; Shen et al. 2021; Yamauchi et al. 2018). However, while limiting library size is ideal if a focussed set of genes is being investigated, this may not be suitable for more exploratory research. One strategy to overcome this is to use a whole-genome *in vivo* screen to identify candidate drug sensitivity genes that are then confirmed by high-throughput validation using a secondary focussed *in vivo* screen with greater statistical power (Yau et al. 2017). Yet, this approach will not resolve false negative hits that are missed in the primary screen and therefore would not be included in the secondary screen, so a high performing whole-genome screen is still preferable. Since our model retained high gRNA representation in early tumours, a short duration whole-genome *in vivo* screen could be used to identify candidate drug sensitivity genes with a permissive FDR threshold. These candidates could then be confirmed in a secondary screen using the same tumour model, but with a focussed gRNA library designed around the candidate genes from the primary screen. Alternatively, a focussed *in vivo* screen could be used to confirm candidate drug sensitivity genes identified in a whole-genome screen in 3D spheroid or organoid cultures (Han et al. 2020; Murakami et al. 2021; Ringel et al. 2020), which would allow gRNA representation to be maintained at a sufficiently high level through the dual screening process, while still accurately modelling many aspects of the human tumour microenvironment.

In summary, we have established a tumour xenograft model with a high frequency of tumour-initiating cells for *in vivo* CRISPR/Cas9 screens. High gRNA representation was observed with small tumours, but there was a considerable loss in representation as tumours grew accompanied by a high degree of random clonal expansion. The major limiting factor in detecting gRNA-driven changes between treatment groups was this clonal expansion which greatly inflates variance in read counts. Based on our findings and binomial thinning simulations, we conclude that our xenograft model would be best suited for short duration *in vivo* screens using a whole-genome gRNA library and/or for longer duration screens using a focussed gRNA library to confirm gene hits originating from a whole-genome *in vivo* or 3D culture screen.

## Authorship note and acknowledgements

The author list on this draft manuscript does not reflect all contributions to this work. We acknowledge the contributions and assistance of Peter Tsai, Cristin Print, Purvi Kakadiya, Stefan Bohlander, Dan Li, Courtney Lynch and Avik Shome to this work.

## Funding acknowledgement

This work was funded by the Auckland Medical Research Foundation, the Marsden Fund of New Zealand and the Health Research Council of New Zealand.

---

**Supp Figure 1:**
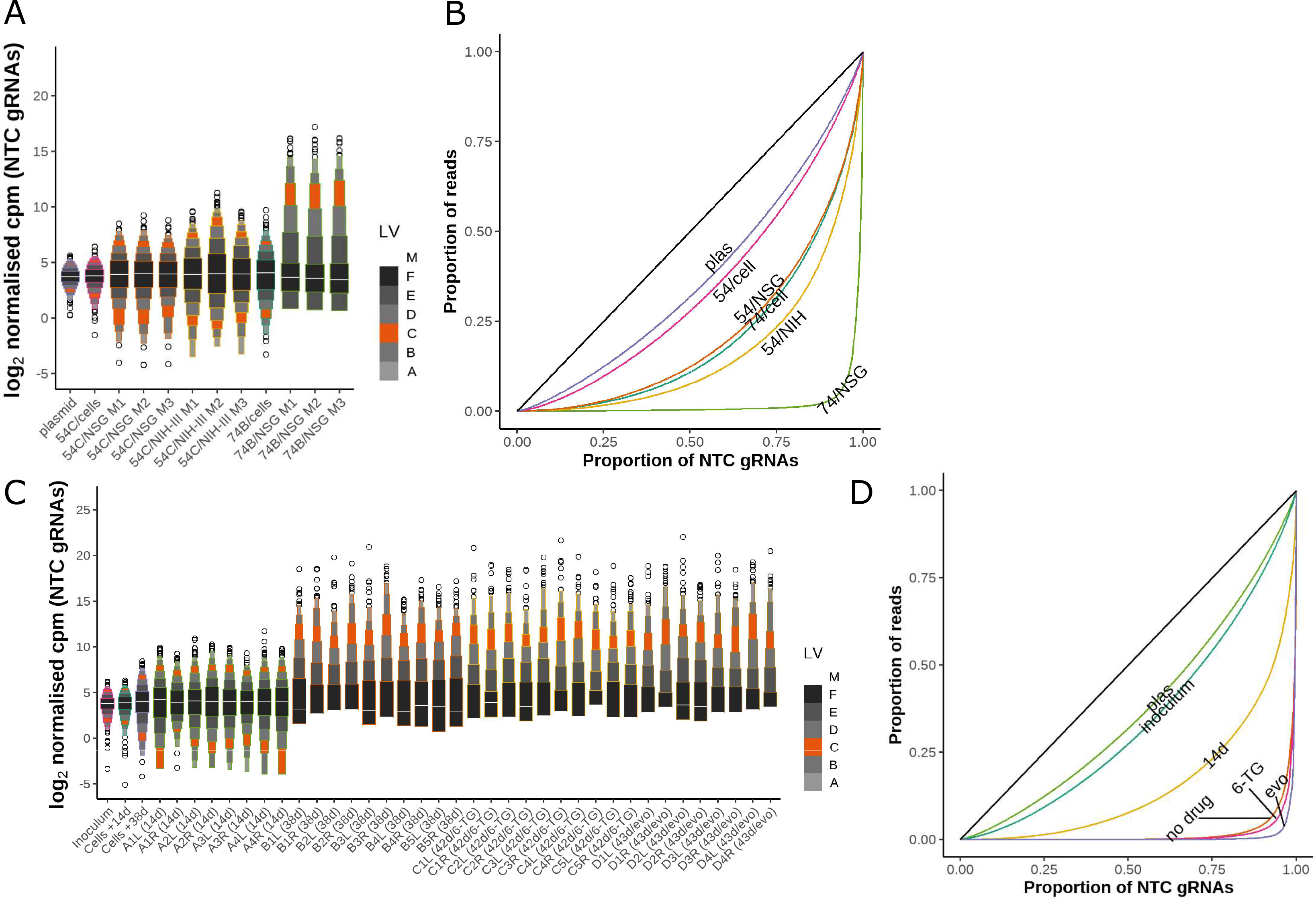
(A) Letter-value plot of log normalised counts per million (log-ncpm) for NTC gRNAs in plasmid, cell inocula and GeCKO tumour samples from the pilot study. (B) Lorenz curves to show distribution of NTC gRNA read counts from the pilot study. (C) Letter-value plot of log normalised counts per million (log-ncpm) for NTC gRNAs in cell inocula and GeCKO tumour samples from the tumour treatment study. (D) Lorenz curves to show distribution of NTC gRNA read counts from the tumour treatment study. For the Lorenz plots, the curve for summed PCR replicates (plasmid and cell samples) or median tumour in a group is highlighted. The black line is the line of equality.

## Notes

### Competing Interest Statement

The authors have declared no competing interest.

